# Curated neurodegeneration axis signals are sensitive to cell composition and analytic choices: an adversarial re-analysis of a transcriptomic scoring pipeline

**DOI:** 10.64898/2026.05.23.727380

**Authors:** Nabanita Ghosh, Krishnendu Sinha

**Affiliations:** Department of Zoology, Maulana Azad College, Kolkata, West Bengal, India; Department of Zoology, Siksha-Bhavana, Visva-Bharati, Santiniketan, West Bengal, India

**Keywords:** Alzheimer’s disease, Parkinson’s disease, gene-set scoring, cell-type composition, competitive null, reproducibility, postmortem brain transcriptomics, analytic flexibility

## Abstract

**Background:** Curated gene-set (“axis”) scores are widely used to summarise disease biology in postmortem brain transcriptomes. In bulk tissue such scores can also change with cell-type composition, with broad transcriptome shifts in diseased tissue, or with analysis choices made after the results were seen. We tested these explanations on our own published pipeline, NeuroFate, whose preprint reported an Alzheimer’s disease (AD) “neuronal vulnerability” axis and a Parkinson’s disease (PD) “divergent” synuclein–mitochondrial axis.

**Results:** A code audit found that the AD discovery axis averaged Sst, Pvalb and Lamp5 interneuron cell-type proportions together with its gene-expression features; that 90 endpoint × axis tests were run before the endpoints were locked; that axes were defined from the cohorts later cited as discovery evidence; and that the PD-divergent label came from an axis-specific rule defined after the result. Their contribution to the AD association could not be quantified because the donor-level data were not archived. In GSE7621 (16 PD, 9 controls) the synuclein– mitochondrial association reproduced (exact *p* = 0.0011) but depended on the scoring method (exact *p* from 0.001 to 0.057; with per-gene standardisation no axis reached Benjamini–Hochberg significance), lay at the 2.4th percentile of 10,000 expression-matched random probe sets (two-sided *p* = 0.06), and was attenuated by 76% (95% interval 26% to *>*100%) after adjustment for a dopaminergic-marker expression proxy (PD coefficient −1.13 to −0.27 SD). A small laser-captured cohort (GSE20141; 10 PD, 8 controls) was inconclusive: no axis reached *p <* 0.05, and the synuclein–mitochondrial effect had the same sign. In 1,000 simulated cohorts, a naive test called a purely composition-driven gene set significant in 82% of cohorts, whereas the proposed battery classified it as composition-independent in 5% while recovering a planted within-cell effect in 98% (moderate effect) and 50% (weak effect) of cohorts.

**Conclusions:** The original axis associations do not establish composition-independent disease programs; their interpretation is sensitive to feature construction, scoring and adjustment for cell composition. Curated axis scores in bulk brain tissue should be reported with competitive null distributions, composition adjustment, exact small-sample inference, and endpoints and scoring fixed before analysis. We provide code for these checks.

## Background

Sample-level gene-set scores are a common way to summarise disease biology in bulk transcriptomes [1–3]. In neurodegeneration research, curated sets labelled “neuroinflammation”, “synuclein–mitochondrial stress” or “neuronal vulnerability” are scored per donor and compared between cases and controls, and a difference is often read as a change in that biological program.

That reading has two well-known weak points. First, a gene-set difference can be non-specific: when much of the transcriptome shifts in diseased tissue, randomly chosen gene sets often separate groups as well as curated ones do [4, 5]. Second, postmortem AD and PD brain loses particular cells—somatostatin-positive and other interneurons early in AD neocortex [6], dopaminergic neurons in PD substantia nigra [7]—so a set that contains cell-type-enriched genes tracks how much of that cell type was sampled rather than what happens inside the cells [8–11]. A third problem is analytic flexibility: choosing endpoints, scoring scales or tests after seeing results inflates false-positive rates even when each choice is defensible on its own [12–14].

We examined all three issues using our own published work as the case. NeuroFate is a command-line package that scores ten curated neurodegeneration axes from donor- or sample-level expression tables [15]. Its preprint reported an AD “neuronal vulnerability” axis discovered in the Seattle Alzheimer’s Disease Brain Cell Atlas (SEA-AD) and nominally replicated in an independent bulk cohort (GSE174367) [16], and a “PD-divergent” synuclein–mitochondrial axis in PD substantia nigra. Here we (i) audit how those results were produced (for the AD result at the level of code, because the derived donor-level data were not archived), (ii) re-test the PD result with exact inference, alternative scoring, a competitive null and composition adjustment, and (iii) measure, by simulation, how often a naive analysis and the proposed set of checks mistake cell loss for disease biology (Fig. 1). Our aim is not a new biomarker but an honest account of what curated axis scores report, and a reusable check for others.

**Figure 1.**
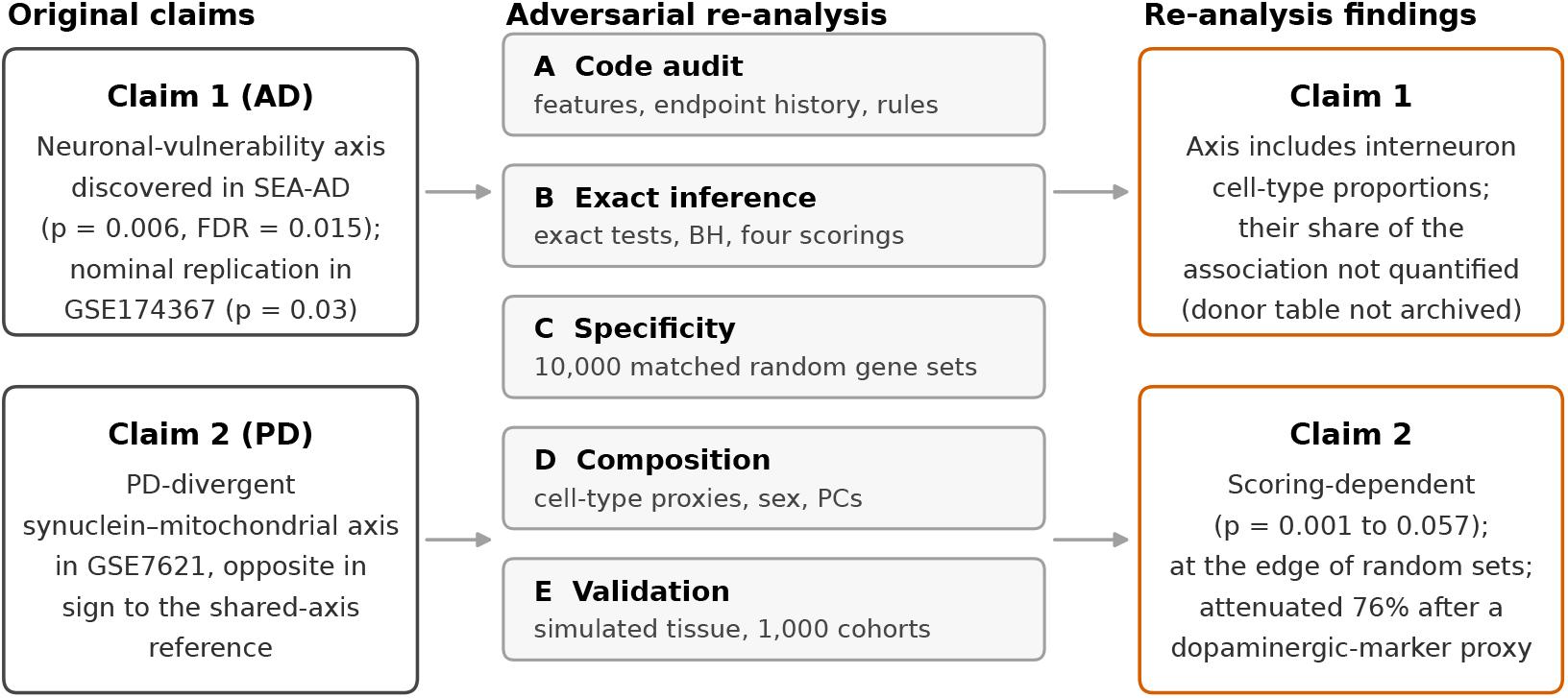
Study overview. The two claims of the original NeuroFate preprint (left) were subjected to five checks (centre): a code audit, exact inference across scoring choices, a competitive null, composition adjustment, and validation of the checks on simulated tissue. The findings for each claim are shown on the right.

## Methods

### Design and disclosure

This is a re-analysis of public data and of the authors’ own analysis code; no new data were generated. The authors wrote the audited software and preprint [15]. Analyses were designed to test the preprint’s claims and were run with code that is archived with every input file and checksum (see Availability of data and materials).

### Original pipeline snapshot

The audit used the NeuroFate analysis code as it stood after the preprint (release candidate 0.3.0; files dated 20 May 2026). The SHA-256 checksum of every audited file matched an independent record made during an internal software audit on 15 October 2026 (Additional file 2, sheet S6). Software issues identified by that audit are listed in Additional file 1, Table S1.

### Code audit

We examined five aspects of how the published results were produced. (1) *Feature construction*. We executed the original feature-matching functions (scripts 22, 106 and 114) on the feature names those scripts generate for SEA-AD. SEA-AD middle temporal gyrus cell types were taken from the consortium’s subclass taxonomy [6]. (2) *Endpoint history*. We compared the association scan of script 107 (phase 21), which selects every metadata column whose name contains a label-like term, with the endpoint registry introduced in script 112 (phase 22). (3) *Classification rules*. We located axis-specific rules and the source of the “reference direction” used to label PD replication (scripts 166 and 171). (4) *Pre-specified directions*. We counted registry entries that accept a change in either direction (“variable” or “… _or_remodeled”). (5) *Statistical implementation*. We compared the original Mann–Whitney implementation (normal approximation without tie or continuity correction) with the exact test.

### Cohorts

GSE7621 comprises postmortem substantia nigra from 16 PD and 9 control donors profiled on Affymetrix HG-U133 Plus 2.0 arrays and processed with MAS5 (global scaling, target intensity 100) [17]. We used the deposited series matrix; its SHA-256 checksum matched the file used by the original analysis. Disease status (“Parkinson’s Disease” or “Old Control”) and sex were taken from the deposited sample characteristics. GSE20141 comprises laser-capture microdissected substantia nigra pars compacta from 10 PD and 8 control donors on the same platform [18]; we re-tested the axis scores that the original pipeline wrote for this cohort. Because rank-based tests are unchanged by monotone transformation of a score, these deposited scores suffice for the association tests reported.

### Axis scores

The ten NeuroFate axes (30 genes) were used as registered. Probes were assigned to genes with the original pipeline’s GPL570 axis-probe map (79 probes, 29 genes; PRKN is not represented). We computed four scores for each axis: (i) the original NeuroFate score (mean of linear-scale probe values per gene, mean across genes, then *z*-score across samples); (ii) the mean of log_2_(*x* + 1) gene values; (iii) the mean of per-gene *z*-scores of log_2_ values (the primary score, which weights genes equally); and (iv) a rank score (the mean within-sample rank of the axis probes among all probes).

### Inference

Effects are rank-biserial correlations (disease minus control). *P*-values are from the exact two-sided Mann– Whitney test [19] and were adjusted across the ten axes with the Benjamini–Hochberg procedure [20]. To reproduce the original result, we also applied the original normal-approximation implementation to the original score. Hedges’ *g* is reported for the primary score.

### Specificity

For each axis, we drew 10,000 random sets of expressed probes (mean log_2_ expression above log_2_ 101; 14,166 probes), one probe for each axis gene, each from the same expression-level bin (20 quantile bins) as the axis gene it replaced, excluding all axis probes [4, 5]. Each random set was scored as the mean of probe *z*-scores, and its rank-biserial effect was computed. We report the observed effect’s percentile and a two-sided empirical *p*-value (the proportion of draws at least as far from the null median as the observed effect, with a +1 correction). Genome-wide annotation was not used, so random sets were drawn at the probe level.

### Composition

A dopaminergic-neuron proxy was defined as the mean *z*-score of six canonical midbrain dopaminergic identity genes [7]: TH (208291_s_at), SLC18A2 (205857_at), DDC (205311_at), ALDH1A1 (212224_at), NR4A2 (204621_s_at) and RET (211421_s_at). A sensitivity proxy added SLC6A3 (206868_at) and KCNJ6 (207446_at). Each axis score was regressed on disease status by ordinary least squares, without and with the proxy, sex, both, or the first two genome-wide principal components of the expressed probes. Attenuation was 1 − *β*_adjusted_*/β*_unadjusted_, with a 95% interval from 2,000 bootstrap resamples drawn within each diagnostic group. We also report Spearman correlations between the axis and the proxy within each group, and the range of exact *p*-values over all leave-one-out subsets.

### Simulation

To measure how often a naive analysis and the proposed checks mistake composition for biology, we simulated bulk tissue from five cell types (neurons, astrocytes, oligodendrocytes, microglia, endothelial cells). Each simulated cohort had 24 disease and 24 control samples. Expression was the sum over cell types of fraction × cell-type profile, multiplied by log-normal per-gene, per-sample variation (SD 0.30 on the natural-log scale). Marker genes for each cell type were the top 100 human BRETIGEA markers [8]; neuronal-vulnerability axis genes were neuron-specific, other axis genes were expressed broadly, and 3,000 background genes had random cell-type loadings. Five scenarios were simulated, 200 cohorts each: no effect; neuron loss only (neuronal fraction ×0.75 in disease); neuron loss plus a direct within-cell increase of inflammatory-microglial axis genes (×1.10, weak, or ×1.20, moderate); and the moderate direct effect alone.

For each cohort and axis we recorded naive significance (exact Mann–Whitney on the per-gene *z*-score, Benjamini–Hochberg across ten axes) and the battery’s classification. An association was classed as *composition-independent* only if it was naively significant, its competitive empirical *p*-value (1,000 expression-matched random gene sets) was below 0.05, and its disease coefficient adjusted for five marker-based cell-type proxies (mean *z*-score of the first 50 BRETIGEA markers per type, excluding axis genes) retained at least half its unadjusted size with a 95% confidence interval excluding zero. Otherwise a significant association was classed as non-specific, composition-explained, or both.

### Chance baselines and power

The probability that six or more of ten axes agree in sign with a reference by chance was taken from the binomial distribution. Power of the two-sided Mann–Whitney test at *α* = 0.05 was estimated from 4,000 simulated normal samples at the sizes of the PD cohorts.

### Software

Analyses used Python 3.13 with NumPy 2.5, pandas 3.0, SciPy 1.18, statsmodels 0.15 and Matplotlib 3.11 [21–25], with a fixed random seed (20261020). A single command (`python run_all.py`) regenerates every number, table and figure from the archived inputs and checks them against the reported values; with the pinned package versions, every output file is byte-identical to the archived one.

### AI-assisted tools

AI-assisted tools were used to support programming and language editing. The authors designed the study, reviewed and verified all code, analyses and text, and take full responsibility for the content of this article.

## Results

### The audit identified four design issues in the original analysis

Table 1 and Fig. 2 summarise the audit. First, the suffix-based feature matcher assigned the SEA-AD cell-type proportions of Sst, Pvalb and Lamp5 interneurons to the neuronal-vulnerability axis, alongside twelve gene-mean and gene-detection features; no other axis received cell-type proportions (Fig. 2A). The discovery axis therefore mixed, with an unknown weight, the abundance of those interneurons with their genes’ expression—and loss of somatostatin-positive interneurons is the earliest cellular change reported by the SEA-AD consortium itself [6]. The gene features are related: SST, PVALB and LAMP5 mark the same interneuron subclasses, so tissue-level means of these genes can also follow composition. Because the features were on different scales and the derived donor-level table was not archived, we could not determine how much each feature type contributed to the score or to its association with dementia; the audit establishes that the score was not a pure expression measure, not that the association is explained by composition.

**Table 1.** Design issues identified in the original analysis.

| Finding | Evidence in the original code or files | Consequence |
| --- | --- | --- |
| Cell-type proportions inside the AD discovery axis | Suffix matcher (scripts 106, 114) assigns `cell_fraction__Sst`, `__Pvalb` and `__Lamp5` to the neuronal-vulnerability axis (15 features on mixed scales); no other axis affected | The SEA-AD association partly measures interneuron abundance, whose early loss SEA-AD itself reported |
| Endpoints locked after an exploratory scan | Script 107 (phase 21) tested 9 SEA-AD endpoints × 10 axes = 90 tests; endpoint registry (script 112, phase 22) kept 4 | Selection after seeing results; reported FDRs do not cover the scan |
| Axes defined from discovery cohorts | Registry cites SEA-AD phases 3–4 and GSE243639 phase 20 as evidence sources | In-sample discovery *p*-values |
| PD-divergent label defined after the result | Script 171 line 65: rule specific to the synuclein–mitochondrial axis; script 166 line 115: reference direction read from observed data (GSE243639 first) | “Divergence” measured against a reference whose own permutation test was not significant |
| Directions that accept either sign | 12 of 20 registry directions accept change in either direction | Directional “consistency” has no pre-registered meaning |
| Small-sample inference | Normal approximation without tie or continuity correction: $p = 0.047$ where exact $p = 0.056$ (5 vs 5) | Nominal significance can be overstated |
| Other | HLA-DRA not matched in two SEA-AD axes; “Mathys 2019” files contain GSE138852 (Grubman et al.) | Gene coverage and file labelling |

**Figure 2.**
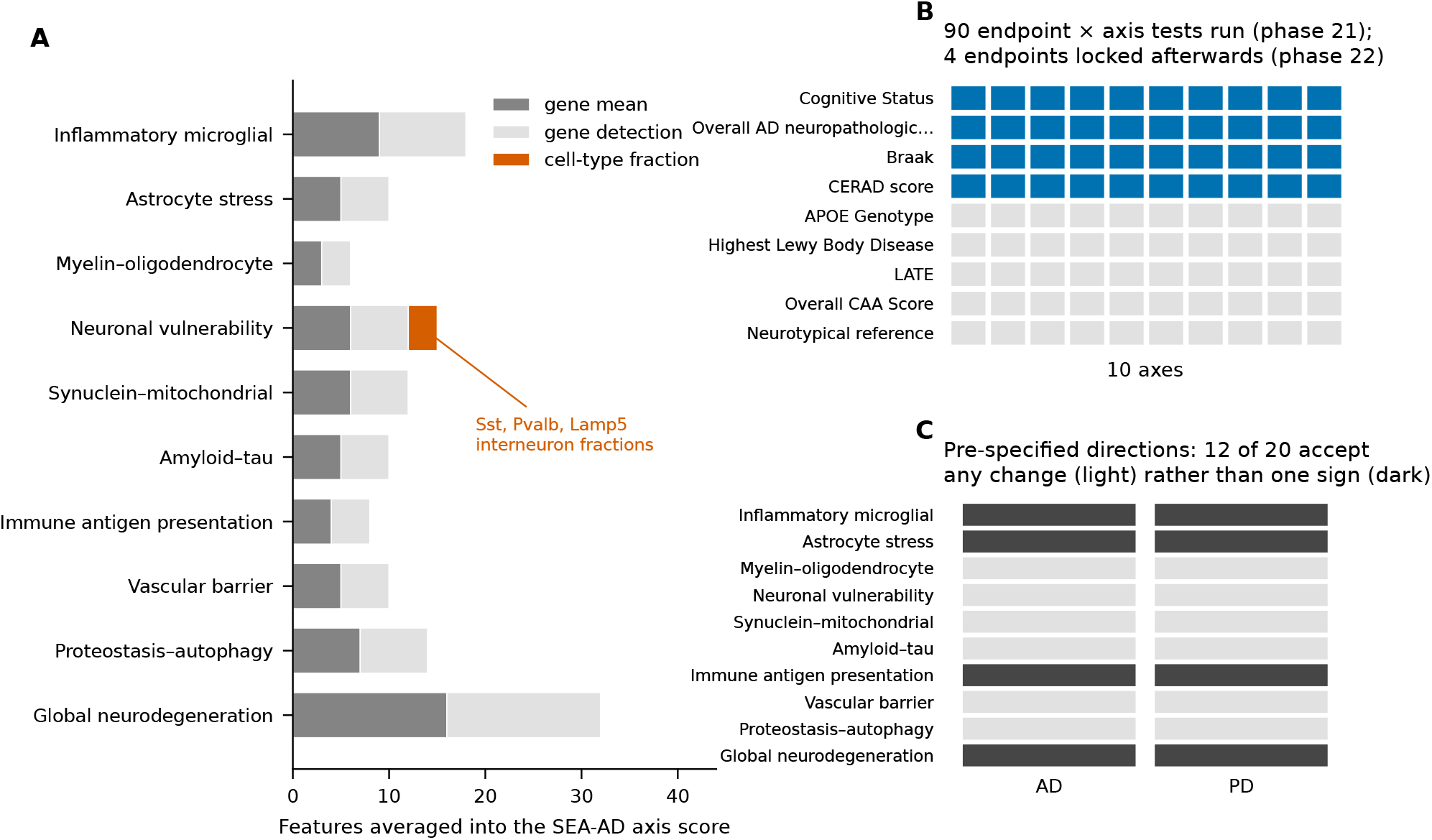
Design audit of the original analysis. (**A**) Features averaged into each SEA-AD axis score by the original matcher, by type; only the neuronal-vulnerability axis received cell-type proportions (Sst, Pvalb and Lamp5 interneurons). (**B**) The phase-21 association scan tested every label-like SEA-AD endpoint against every axis (90 tests); four endpoints were locked afterwards. (**C**) Pre-specified directions in the axis registry: light cells accept a change in either direction.

Second, the association scan in phase 21 tested every label-like SEA-AD column against every axis (9 endpoints × 10 axes = 90 tests, including APOE genotype, LATE, cerebral amyloid angiopathy and Lewy body stage) before the endpoint registry that locked four endpoints was introduced in phase 22 (Fig. 2B). The reported false-discovery rates do not account for this scan. Third, the axis registry names SEA-AD and GSE243639 analyses as each axis’s evidence source, so axes were assembled after those cohorts had been examined and the same cohorts were then cited as discovery evidence.

Fourth, the “PD-divergent” label was produced by a rule written for one axis: script 171 contains a branch that applies only when the synuclein–mitochondrial axis is negative, significant and opposite to its reference. That reference direction was itself read from observed data, first from GSE243639, a cohort whose own permutation test was not significant (*p* = 0.109). Twelve of the twenty pre-specified directions in the registry accept a change in either direction (Fig. 2C), so directional “consistency” had no pre-registered meaning for most axes. With ten axes and no true effects, six or more are directionally consistent with any reference 38% of the time.

The original Mann–Whitney implementation also used a normal approximation that is inexact at small sample sizes: for two groups of five, it returned *p* = 0.047 where the exact test gives *p* = 0.056. Two smaller issues were found: HLA-DRA was not matched in two SEA-AD axes because of a hyphen–underscore difference, and files labelled “Mathys 2019” in the repository contain GSE138852, a different study [26, 27]. Software issues identified by the internal audit are listed for completeness in Additional file 1, Table S1.

### GSE7621: the synuclein–mitochondrial association depends on how it is scored

The original result reproduced exactly: with the original score and test, the synuclein–mitochondrial axis was lower in PD (rank-biserial −0.76, *p* = 0.0018). The exact test gave *p* = 0.0011, so the original association does not depend on the normal approximation. On the original linear scale, however, 48% of this axis came from APOE, an astrocyte-enriched gene; other axes were similarly dominated by single genes (GFAP 70% of the astrocyte-stress axis, B2M 88% of the vascular-barrier axis). With per-gene standardisation, which weights genes equally, the effect was −0.60 (exact *p* = 0.014; Benjamini–Hochberg-adjusted 0.14), and no axis reached adjusted significance (Fig. 3A; Table 2). Across the four scores the exact *p*-value for this axis ranged from 0.001 to 0.057 (Fig. 3B), and which axes reached *p <* 0.05 depended on the score: synuclein–mitochondrial under three scores, amyloid–tau under three, myelin–oligodendrocyte under one (Additional file 1, Fig. S2). Leave-one-out *p*-values for the primary score ranged from 0.002 to 0.032.

**Table 2.** The GSE7621 synuclein–mitochondrial association under alternative analyses. (16 PD, 9 control). OLS coefficients are in SD units of the per-gene *z*-score axis, with 95% confidence intervals.

| Analysis | Effect | $p$-value |
| --- | --- | --- |
| Original score, original test (normal approximation) | rank-biserial $-0.76$ | 0.0018 |
| Original score, exact test | $-0.76$ | 0.0011 |
| $\log_2$ mean, exact test | $-0.61$ | 0.012 |
| Per-gene z-score (primary), exact test | $-0.60$ | 0.014 (BH-adjusted 0.14) |
| Rank score, exact test | $-0.47$ | 0.057 |
| Competitive null, 10,000 matched random probe sets | 2.4th percentile | 0.062 (two-sided empirical) |
| OLS: PD only | $-1.13$ SD ($-1.86$ to $-0.40$) | 0.004 |
| OLS: + sex | $-1.26$ SD ($-2.06$ to $-0.47$) | 0.003 |
| OLS: + dopaminergic proxy | $-0.27$ SD ($-1.09$ to $0.56$) | 0.51 |
| OLS: + dopaminergic proxy + sex | $-0.40$ SD ($-1.24$ to $0.44$) | 0.33 |
| OLS: + dopaminergic proxy (8 probes) | $-0.38$ SD ($-1.32$ to $0.55$) | 0.41 |
| OLS: + PC1, PC2 | $-0.85$ SD ($-1.66$ to $-0.05$) | 0.039 |
| Attenuation by dopaminergic proxy | 76% (bootstrap 26% to $>100\%$) | – |
| Leave-one-out range, primary score | – | 0.002 to 0.032 |

**Figure 3.**
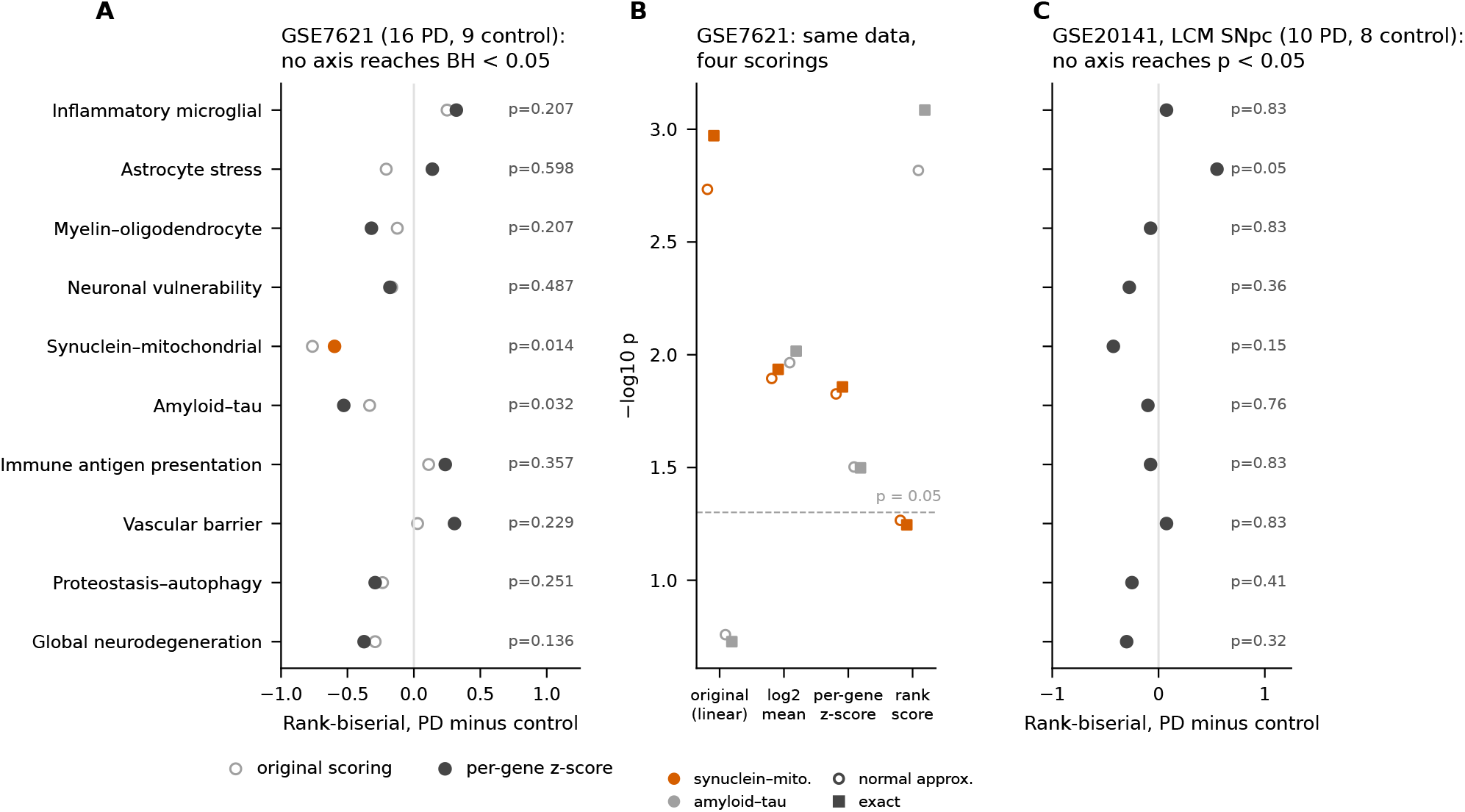
Association tests in PD substantia nigra. (**A**) GSE7621: rank-biserial effect (PD minus control) for each axis with the per-gene *z*-score (filled; exact *p* shown) and the original linear score (open). (**B**) GSE7621: *p*-values of the synuclein–mitochondrial and amyloid–tau axes under four scorings and two tests; dashed line, *p* = 0.05. (**C**) GSE20141 (laser-captured SNpc): axis effects recomputed from the original pipeline’s deposited scores, with exact *p*-values.

At the gene level, the decrease was carried by PINK1 (rank-biserial −0.63, *p* = 0.010), SNCA (−0.53, *p* = 0.032) and LRRK2 (−0.44, *p* = 0.074); APOE did not differ (*p* = 0.15). The amyloid–tau axis (primary *p* = 0.032, adjusted 0.16) was driven mainly by PSEN2 (−0.78, *p* = 0.0008).

### GSE7621: the association lies at the edge of the range of matched random probe sets

PD substantia nigra showed a broad transcriptional shift: 17.2% of 14,166 expressed probes had *p <* 0.05, against 5% expected by chance. Against 10,000 expression-matched random probe sets, the synuclein–mitochondrial effect sat at the 2.4th percentile (two-sided empirical *p* = 0.062) and the amyloid–tau effect at the 4.0th percentile (*p* = 0.092); every other axis lay well inside the random range (Fig. 4). The synuclein–mitochondrial axis is therefore associated with PD, but its association is not clearly exceptional among comparably expressed probe sets in this tissue; a non-significant competitive test does not show that the axis behaves like a random set.

**Figure 4.**
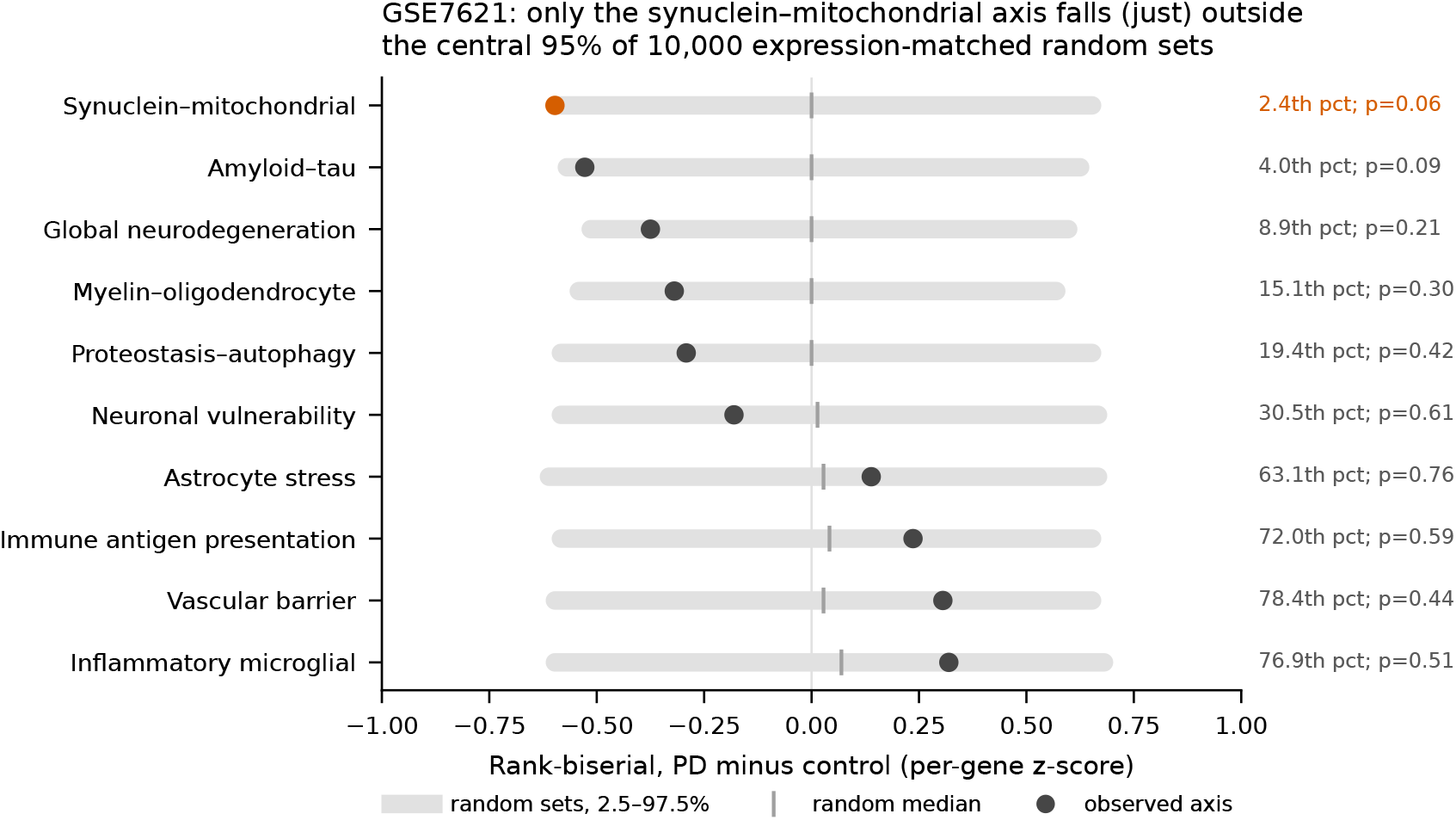
Competitive null in GSE7621. For each axis, the observed effect (dot) is compared with the central 95% (grey bar) and median (tick) of 10,000 random sets of expressed probes matched to the axis genes on expression level. Right: observed percentile and two-sided empirical *p*-value.

### GSE7621: the association is attenuated after adjustment for a dopaminergic-marker proxy

The six dopaminergic markers were tightly co-expressed (median pairwise Spearman *ρ* = 0.77) and strongly reduced in PD (log_2_ fold changes −1.1 to −3.5; proxy rank-biserial −0.78, *p* = 0.0008; Fig. 5C). The synuclein– mitochondrial axis tracked this proxy (*ρ* = 0.56 overall; 0.47 within PD and 0.62 within controls; Fig. 5A). Adjusting for the proxy reduced the PD coefficient from −1.13 SD (95% CI −1.86 to −0.40; *p* = 0.004) to −0.27 SD (−1.09 to 0.56; *p* = 0.51), an attenuation of 76% (bootstrap 95% interval 26% to more than 100%), and raised *R*^*2*^ from 0.31 to 0.54 (Fig. 5B; Table 2). The result was similar with sex added (−0.40 SD, *p* = 0.33) or with the eight-probe proxy (−0.38 SD, *p* = 0.41). Two genome-wide principal components, which carry composition and other variation, attenuated the effect less (− 0.85 SD, *p* = 0.039). Adjustment for sex alone did not attenuate it; the PD group was mostly male (19% female, against 56% of controls; Fisher *p* = 0.087). The amyloid–tau axis was not attenuated by the dopaminergic proxy (−1.05 to −1.23 SD). Because the proxy is itself an expression measure and dopaminergic neuron loss is part of PD, these models estimate the association conditional on the proxy: they do not show that 76% of the effect is caused by neuron loss, and they may remove disease effects that act through that loss.

**Figure 5.**
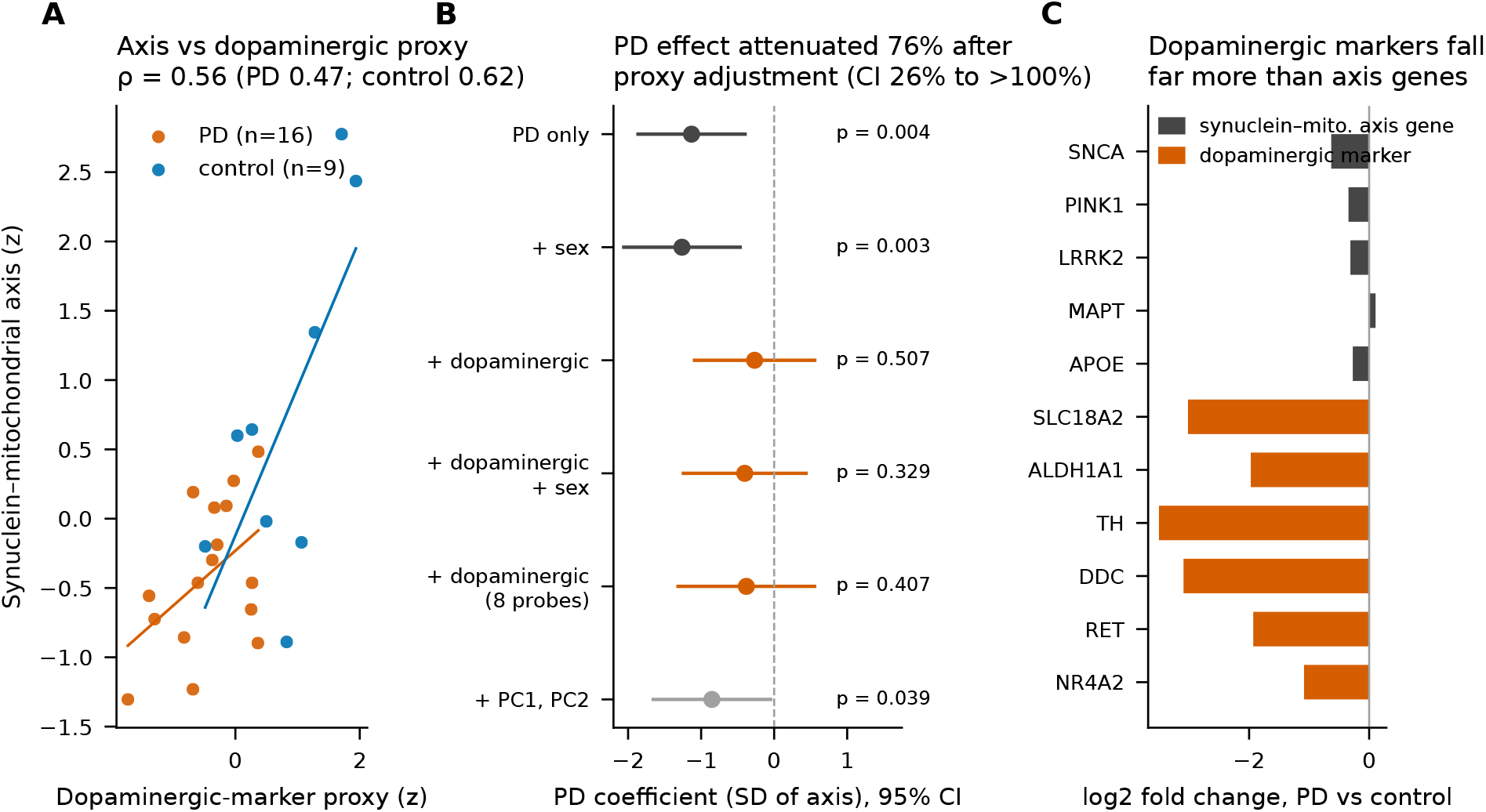
Composition in GSE7621. (**A**) The synuclein–mitochondrial axis against the dopaminergic-marker proxy, with within-group least-squares lines. (**B**) PD coefficient (SD of the axis) with 95% CI under alternative adjustments; the title gives the attenuation by the dopaminergic proxy with its bootstrap interval. (**C**) log_2_ fold change (PD vs control) of the synuclein–mitochondrial axis genes and of the dopaminergic markers.

### GSE20141: an inconclusive small-cohort replication

In laser-captured substantia nigra pars compacta, which is enriched for neurons by design, no axis reached *p <* 0.05 (smallest *p* = 0.055, astrocyte stress; Fig. 3C). The synuclein–mitochondrial axis was lower in PD (− 0.43, *p* = 0.15), in the same direction as in GSE7621; the “divergence” of GSE7621 was therefore relative only to GSE243639. The ten nominally distinct axes were strongly inter-correlated (median Spearman *ρ* = 0.61), and per-sample axis means on the original linear scale varied 7.8-fold (Additional file 1, Fig. S1). With 10 and 8 samples, power to detect even a standardised difference of 1.0 was 0.44 (Additional file 1, Fig. S6), so this cohort neither confirms nor excludes an effect of the size seen in GSE7621.

### Simulation: naive axis tests mistake cell loss for biology

In simulated tissue with neuron loss but no within-cell change, the naive test called the neuron-specific gene set significant in 82% of cohorts, whereas the battery classified it as composition-independent in 5.0% (Fig. 6A). With concurrent neuron loss, a planted within-cell change in the inflammatory gene set was classified as composition-independent in 98% of cohorts at the moderate effect size and 50% at the weak effect size, and in 100% when there was no composition change (Fig. 6B). With no effect at all, 0.8% of axis tests were classed as composition-independent (6% of cohorts had at least one such call among ten axes). Gene sets that shared genes with the planted set were also detected, as expected (Additional file 1, Fig. S5).

**Figure 6.**
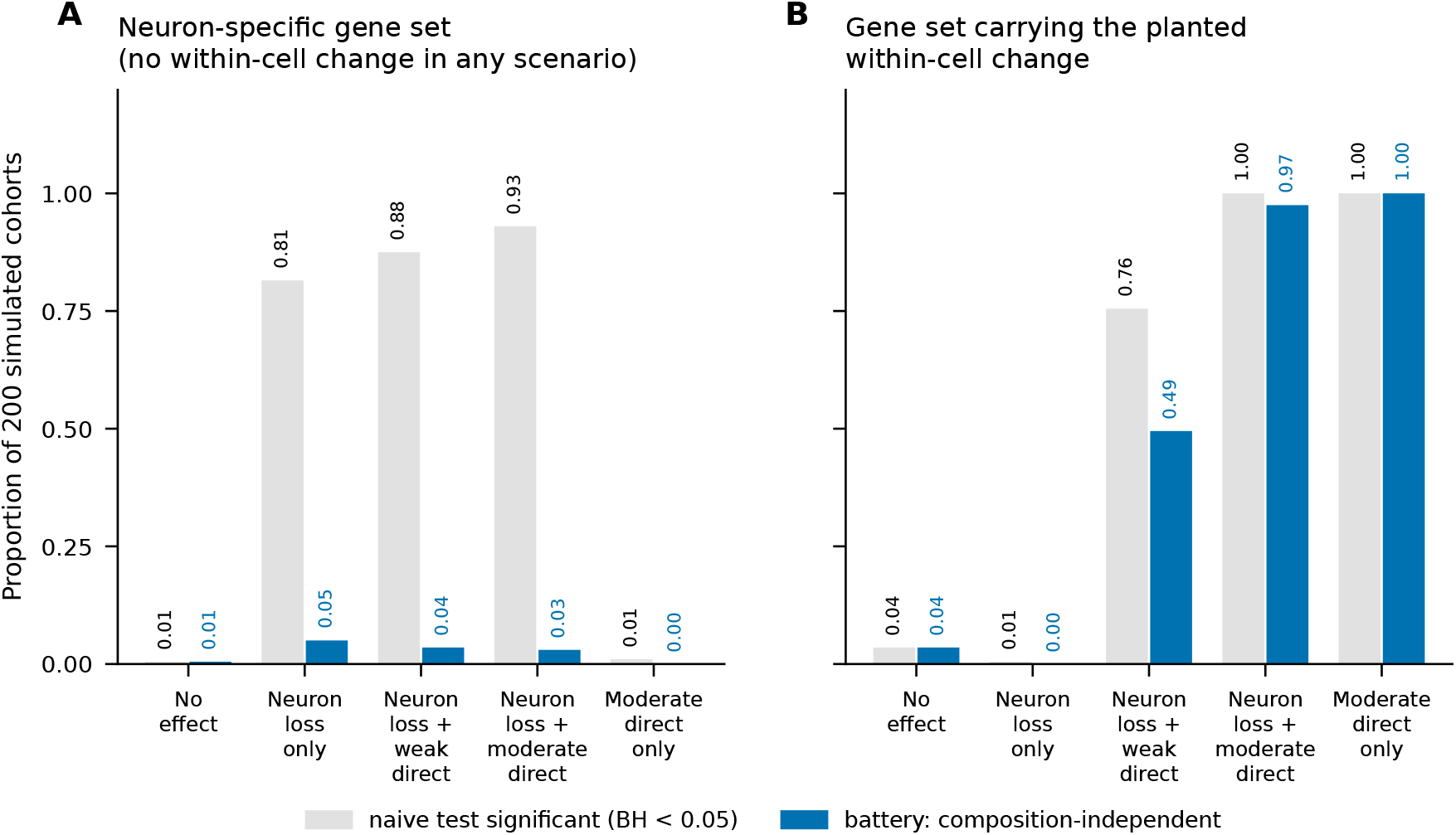
Operating characteristics on simulated tissue. Proportion of 200 simulated cohorts per scenario in which the naive test was significant (grey) and the battery classified the association as composition-independent (blue), for (**A**) a neuron-specific gene set that never changes within cells, and (**B**) the gene set carrying a planted within-cell change.

## Discussion

The re-analysis does not support reading either original association as evidence of a composition-independent disease program, but it also does not show that the underlying associations are absent. The AD neuronal-vulnerability axis averaged interneuron proportions into its score, and its gene features mark the same interneurons; its association with dementia may therefore partly reflect the early interneuron loss reported in SEA-AD [6], but how much it does could not be measured here. The PD synuclein–mitochondrial association reproduced in GSE7621—SNCA- and PINK1-containing scores are lower in PD substantia nigra—but its strength depended on a scoring choice that let APOE dominate, it lay at the edge of the range of matched random probe sets in a tissue where much of the transcriptome shifts, and most of it was attenuated after adjustment for a dopaminergic-marker expression proxy. A lower SNCA signal in nigra that has lost dopaminergic neurons is expected [7]; on its own, it does not indicate a distinct, divergent program.

The mechanisms are general. A score built from cell-type-enriched genes inherits the composition of the sample [8–11]. Averaging on a linear scale lets one abundant gene dominate. In diseased tissue a curated set must be compared with random sets of similar expression, not with zero [4, 5]. Analytic flexibility can also enter a pipeline designed to prevent it: here the endpoints were locked after an exploratory scan, and a classification rule was defined after the result was known [12–14]. Each step was reasonable in isolation, which is why such issues are easy to miss. Our simulation shows the size of the risk: under the noise level we simulated, a naive test reported a composition-only gene set as significant in four of five cohorts.

### Recommendations

Before interpreting a curated gene-set score from bulk brain tissue as biology, we suggest that analysts:

1. score on a log or rank scale with per-gene standardisation, and report how results change across reasonable scorings;
2. compare each effect with expression-matched random gene sets;
3. adjust for cell-type proxies built from an independent marker resource, and report attenuation with uncertainty;
4. use exact or permutation inference at small sample sizes, with multiplicity stated over everything tested;
5. fix endpoints, scoring and decision rules before analysing the data, and report analyses done earlier as exploratory;
6. where possible, confirm effects in a neuron-enriched or single-nucleus design.

### Limitations

The AD claim was audited at the level of code only. The derived SEA-AD donor-level feature table, the cell-level expression from which it was built and the GSE174367 data were not archived with the original analysis code, so we could not reproduce the original AD association, compare it with expression-only and composition-only scores, adjust it for measured cell fractions, test expression within interneuron subclasses, or re-analyse the GSE174367 replication (*p* = 0.03, false-discovery rate 0.24 as originally reported). The AD association may therefore retain a component that is independent of composition. For GSE20141 we re-tested the scores deposited by the original pipeline rather than recomputing them from the expression data. Our dopaminergic proxy uses six marker probes; marker proxies are imperfect, and adjusting for cell loss can remove true disease effects that act through that loss, so we report attenuation rather than proof of absence. With 25 samples, the adjusted GSE7621 model cannot exclude a moderate residual effect (95% CI −1.09 to 0.56 SD), and the attenuation interval is wide. The competitive null was drawn at the probe level and matched on expression only, because genome-wide gene annotation was not used; it does not account for gene-gene correlation or cell-type specificity. The simulation is stylised; it shows how the checks behave, not the composition of real tissue. Finally, the authors developed the audited software and analysed these cohorts before; we have disclosed this and archived every step. Independent confirmation of the battery in larger AD and PD cohorts, under a plan fixed in advance, is the natural next step.

## Conclusions

Curated neurodegeneration axis scores computed from postmortem bulk brain tissue can reflect cell composition, scoring choices and analytic flexibility as well as disease-related expression changes. In our own pipeline, the AD axis mixed interneuron proportions with expression features, and the PD-divergent association was scoring-dependent, at the edge of the range of matched random probe sets and largely attenuated after adjustment for a dopaminergic-marker proxy. The original associations therefore do not establish composition-independent disease programs. Competitive nulls, composition adjustment, exact inference and pre-specified analysis plans help separate these explanations, and the code we provide applies them.

## Availability and requirements

- **Project name:** NeuroFate composition audit (reproducibility package for this article)
- **Project home page:** https://doi.org/10.5281/zenodo.23491606
- **Operating system(s):** platform independent
- **Programming language:** Python (≥ 3.12)
- **Other requirements:** NumPy, pandas, SciPy, statsmodels, Matplotlib, openpyxl (versions pinned in the package)
- **License:** MIT (code); CC BY 4.0 (derived data, figures and documentation)
- **Any restrictions to use by non-academics:** none

### Abbreviations

AD: Alzheimer’s disease
BH: Benjamini–Hochberg
CI: confidence interval
LCM: laser-capture microdis-section
MAS5: Affymetrix Microarray Suite 5.0
OLS: ordinary least squares
PC: principal component
PD: Parkinson’s disease
SD: standard deviation
SEA-AD: Seattle Alzheimer’s Disease Brain Cell Atlas
SNpc: substantia nigra pars compacta

## Declarations

### Ethics approval and consent to participate

Not applicable. This study re-analysed publicly available, de-identified data that were collected under the original studies’ ethical approvals.

### Consent for publication

Not applicable.

### Availability of data and materials

The datasets analysed are available in the Gene Expression Omnibus under accessions GSE7621 and GSE20141. The audited NeuroFate code is available at https://github.com/sinhakrishnendu/NeuroFate. All code, input files with SHA-256 checksums, derived results, figures and the simulation are archived at Zenodo (doi:10.5281/zenodo.23491606) [28] and reproduce every number in this article with a single command. All numerical results are also provided in Additional file 2.

### Authors’ contributions

NG and KS conceived the re-analysis. KS performed the code audit, re-analyses and simulation and prepared the reproducibility package. NG and KS interpreted the results. NG and KS wrote and revised the manuscript. Both authors read and approved the final manuscript.

## Acknowledgements

We thank the investigators who generated and deposited the datasets analysed here.

## Additional files

**Additional file 1** (PDF). Supplementary information: Supplementary Methods; Table S1 (software issues identified by the internal audit); Tables S2–S8 (feature matching, GSE20141 and GSE7621 results, dopaminergic probes, simulation summary); Figures S1–S6 (GSE20141 axis correlations, GSE7621 specification grid, null distributions, leave-one-out stability, simulation class distributions, power).

**Additional file 2** (XLSX). All numerical results, one sheet per analysis (sheets S1–S17), generated by the archived code.

