## Additional file 1 for "Curated neurodegeneration axis signals are sensitive to cell composition and analytic choices: an adversarial re-analysis of a transcriptomic scoring pipeline"

Contents: Supplementary Methods S1–S5; Tables S1–S8; Figures S1–S6. All values are generated by the archived code; the same numbers are in Additional file 2.

### Supplementary Methods

#### S1. Audited files

The audit executed or inspected these files from the original NeuroFate repository (release candidate 0.3.0, dated 20 May 2026): scripts `22_build_donor_feature_table.py`, `106_build_neurofate_axis_scores.py`, `107_test_neurofate_axis_associations.py`, `112_test_axis_associations_endpoint_locked.py`, `114_run_endpoint_locked_random_axis_controls.py`, `166_test_phase34_pd_axis_replication.py` and `171_audit_gse7621_axis_direction_and_probe_mapping.py`; registries `neurofate_axis_registry.tsv`, `neurofate_axis_endpoint_registry.tsv` and `target_gene_panel_v1.tsv`; and the GSE20141 outputs `axis_scores.tsv` and `gpl570_axis_probe_mapping.tsv`. SHA-256 checksums of all scripts and both registries matched the record of an internal audit made on 15 October 2026 (Additional file 2, sheet S6). The target-gene panel was not in that record; its checksum is listed in the package manifest.

#### S2. Reproducing the SEA-AD feature names

Script 22 names donor-level features `gene_mean__<gene>`, `gene_detection__<gene>`, `index__<index>` and `cell_fraction__<Subclass>`, after replacing non-alphanumeric characters with underscores. We generated these names for the 30-gene target panel, the six donor indices defined in script 22, and the 24 SEA-AD middle temporal gyrus subclasses (Astrocyte, Chandelier, Endothelial, L2/3 IT, L4 IT, L5 ET, L5 IT, L5/6 NP, L6 CT, L6 IT, L6 IT Car3, L6b, Lamp5, Lamp5 Lhx6, Microglia-PVM, OPC, Oligodendrocyte, Pax6, Pvalb, Sncg, Sst, Sst Chodl, VLMC, Vip). We then called the original `axis_feature_map` function, which assigns a feature to an axis when the feature name ends in `_<GENE>` or contains `__<GENE>__` (case-insensitive). Because Lamp5, Pvalb and Sst are both subclass names and gene symbols in the neuronal-vulnerability axis, their cell-type fractions are assigned to it. The exact subclass labels in the original donor table should be confirmed against that table; the matching rule itself is reproduced from the original code.

#### S3. Original Mann–Whitney implementation

Scripts 112 and 166 compute $U$ from mid-ranks and return

$$p = 2\left[1-\Phi(|z|)\right], \qquad z = \frac{U - n_1 n_2/2}{\sqrt{n_1 n_2 (n_1+n_2+1)/12}},$$

without tie correction, continuity correction or an exact distribution. For $[1,2,3,4,8]$ vs $[5,6,7,9,10]$ this gives $p = 0.0472$; the exact two-sided test gives $p = 0.0556$.

#### S4. Competitive null in GSE7621

Expressed probes were those with mean $\log_2(x+1)$ above $\log_2 101$ (14,166 of 54,339 probes, AFFX controls excluded). Probes were divided into 20 equal-count bins by mean expression. For each axis gene, the bin was the median bin of its probes. Each random set drew one probe per axis gene from that gene's bin, excluding all 79 axis probes, with replacement across draws. The random set's score was the mean of its probes' $z$-scores;

its rank-biserial effect was computed with the asymptotic Mann–Whitney statistic. The empirical two-sided $p$-value was $(1+\#\{|\text{draw}-\text{median}| \geq |\text{observed}-\text{median}|\})/(10{,}000+1)$.

#### S5. Simulation

**Simulation parameters.**

| Parameter | Value |
|---|---|
| Cohorts per scenario | 200 (24 disease, 24 control samples each) |
| Cell types and base fractions | neurons 0.40, astrocytes 0.20, oligodendrocytes 0.25, microglia 0.08, endothelial 0.07; per-sample log-normal jitter (SD 0.12), renormalised |
| Marker genes | first 100 BRETIGEA human markers per cell type (excluding axis genes), loading 1.0 on own type and 0.05 on others |
| Axis genes | neuronal-vulnerability genes neuron-specific (1.0 vs 0.02); all other axis genes 0.3 on every type |
| Background genes | 3,000 genes with Dirichlet$(1,\ldots,1)$ loadings |
| Gene scale | log-normal(6, 1.2) per gene |
| Noise | log-normal(0, 0.30) per gene and sample |
| Scenarios | S0 no effect; S1 neuronal fraction $\times 0.75$ in disease; S2 S1 plus inflammatory-microglial axis genes $\times 1.10$ (weak) or $\times 1.20$ (moderate) within cells; S3 $\times 1.20$ without composition change |
| Battery | exact Mann–Whitney on per-gene $z$-score with BH over 10 axes; 1,000 expression-matched random gene sets; OLS adjusted for five proxies (first 50 BRETIGEA markers per type) |
| Seed | 20261020 |

Every analysis can be reproduced with one command (`python run_all.py`) from the archived package (Zenodo, doi:10.5281/zenodo.23491606); see its README for the mapping of scripts to tables and figures.

#### Supplementary Tables

**Table S1. Software issues identified by the internal audit of 15 October 2026.** These issues concern the NeuroFate software, were identified with targeted test inputs, and are listed for completeness.

| ID | Issue | Location | Effect observed |
|---|---|---|---|
| F01 | Explicit endpoint classes overridden by built-in aliases | `axis.py:51` | AD counted as positive when the user set AD as the negative class; PD counted in an AD-vs-control contrast |
| F02 | Numeric endpoint column treated as an axis | `axis.py:284` | Samples with identical expression scored 0.5 and 1.0 because their labels differed |
| F03 | Mixed exact and normalised sample matches | `ingest.py:739` | Join report 2/2 matched, but one sample not retained |
| F04 | Sample-identifier normalisation collisions | `ingest.py:139, 743` | Data assigned to the wrong sample |
| F05 | Conflicting duplicate metadata | `ingest.py:653` | Endpoint label determined by row order |
| F06 | Benchmark split ignores donor identity | `benchmarking.py:146` | Random labels: AUROC 0.375; same donors repeated ten times: AUROC 1.0 |
| F07 | Permutation control does not permute the actual target | `script 59; benchmarking.py:75` | Original disease signal retained under the null |
| F08 | Hand-written rank test without exact or tie handling | `scripts 112, 166` | $p = 0.047$ where exact $p = 0.056$ |
| F09 | Package build configuration | `pyproject.toml` | Superseded licence classifier rejected by setuptools $\geq 77$ |
| F10 | Empty feature list restores all features | `benchmarking.py:155` | Ablation of every feature still used the features |
| F11 | Non-finite or missing values not flagged | `axis.py:208, 295` | Infinite risk score reported as a successful run with full coverage |
| F12 | Endpoint and sample-column inference depends on names and order | `ingest.py:280, 296` | Swapping column order changed the selected endpoint |
| F13 | Unknown genotype labels | `benchmarking.py:72` | Missing APOE treated as negative, or crash |
| F14 | Test suite not self-contained | `tests/` | 39 of 454 tests did not pass in a clean environment; version strings inconsistent |

**Table S2. Features assigned to each SEA-AD axis by the original matcher.**

| Axis | Features | Cell-type fractions | Genes not matched | Registered evidence source |
|---|---|---|---|---|
| Inflammatory microglial | 18 | – | – | preliminary_sea_ad_phase3_4_and_gse243639_phase20 |
| Astrocyte stress | 10 | – | – | preliminary_sea_ad_phase3_4_and_gse243639_phase20 |
| Myelin–oligodendrocyte | 6 | – | – | preliminary_sea_ad_phase3_4_and_gse243639_phase20 |
| Neuronal vulnerability | 15 | Lamp5; Pvalb; Sst | – | preliminary_sea_ad_phase3_4_and_gse243639_phase20 |
| Synuclein–mitochondrial | 12 | – | – | gse243639_phase20_primary_pd_axis |
| Amyloid–tau | 10 | – | – | sea_ad_primary_ad_axis |
| Immune antigen presentation | 8 | – | HLA-DRA | preliminary_sea_ad_phase3_4_and_gse243639_phase20 |
| Vascular barrier | 10 | – | – | limited_target_panel_support |
| Proteostasis–autophagy | 14 | – | – | preliminary_sea_ad_phase3_4_and_gse243639_phase20 |
| Global neurodegeneration | 32 | – | HLA-DRA | integrative_axis_across_sea_ad_and_gse243639 |

**Table S3. GSE20141 axis tests** (10 PD, 8 control; recomputed from the original pipeline's deposited axis scores).

| Axis | Rank-biserial | $p$ (original method) | $p$ (exact) | BH |
|---|---|---|---|---|
| Inflammatory microglial | 0.07 | 0.790 | 0.829 | 0.829 |
| Astrocyte stress | 0.55 | 0.051 | 0.055 | 0.545 |
| Myelin–oligodendrocyte | −0.07 | 0.790 | 0.829 | 0.829 |
| Neuronal vulnerability | −0.28 | 0.328 | 0.360 | 0.816 |
| Synuclein–mitochondrial | −0.43 | 0.131 | 0.146 | 0.729 |
| Amyloid–tau | −0.10 | 0.722 | 0.762 | 0.829 |
| Immune antigen presentation | −0.07 | 0.790 | 0.829 | 0.829 |
| Vascular barrier | 0.07 | 0.790 | 0.829 | 0.829 |
| Proteostasis–autophagy | −0.25 | 0.374 | 0.408 | 0.816 |
| Global neurodegeneration | −0.30 | 0.286 | 0.315 | 0.816 |

**Table S4. GSE7621 axis tests** (16 PD, 9 control). Primary: per-gene $z$-score, exact test, BH over 10 axes. Original: linear score, normal approximation.

| Axis | Genes | Rank-biserial | $p$ (exact) | BH | Hedges' $g$ | Original rank-biserial | Original $p$ | Largest gene share (linear) |
|---|---|---|---|---|---|---|---|---|
| Inflammatory microglial | 9 | 0.32 | 0.207 | 0.359 | 0.50 | 0.25 | 0.308 | CX3CR1 20% |
| Astrocyte stress | 5 | 0.14 | 0.598 | 0.598 | −0.04 | −0.21 | 0.396 | GFAP 70% |
| Myelin–oligodendrocyte | 3 | −0.32 | 0.207 | 0.359 | −0.49 | −0.12 | 0.610 | PLP1 64% |
| Neuronal vulnerability | 6 | −0.18 | 0.487 | 0.541 | −0.25 | −0.17 | 0.497 | NEFL 45% |
| Synuclein–mitochondrial | 5 | −0.60 | 0.014 | 0.139 | −1.29 | −0.76 | 0.002 | APOE 48% |
| Amyloid–tau | 5 | −0.53 | 0.032 | 0.159 | −1.16 | −0.33 | 0.174 | APOE 44% |
| Immune antigen presentation | 5 | 0.24 | 0.357 | 0.447 | 0.40 | 0.11 | 0.651 | B2M 78% |
| Vascular barrier | 5 | 0.31 | 0.229 | 0.359 | 0.34 | 0.03 | 0.910 | B2M 88% |
| Proteostasis–autophagy | 6 | −0.29 | 0.251 | 0.359 | −0.64 | −0.24 | 0.336 | APP 41% |
| Global neurodegeneration | 16 | −0.38 | 0.136 | 0.359 | −0.55 | −0.29 | 0.234 | PLP1 36% |

**Table S5. GSE7621 gene-level differences** ($\log_2$; exact Mann–Whitney).

| Gene (probe) | Role | $\log_2$ FC (PD − control) | Rank-biserial | $p$ (exact) |
|---|---|---|---|---|
| SLC18A2 (205857_at) | dopaminergic proxy | −3.02 | −0.86 | $1.4 \times 10^{-4}$ |
| ALDH1A1 (212224_at) | dopaminergic proxy | −1.97 | −0.85 | $1.9 \times 10^{-4}$ |
| PSEN2 | Amyloid–tau | −0.60 | −0.78 | $8.2 \times 10^{-4}$ |
| TH (208291_s_at) | dopaminergic proxy | −3.50 | −0.75 | 0.001 |
| DDC (205311_at) | dopaminergic proxy | −3.09 | −0.71 | 0.003 |
| RET (211421_s_at) | dopaminergic proxy | −1.93 | −0.69 | 0.004 |
| PINK1 | Synuclein–mitochondrial; Proteostasis–autophagy; Global neurodegeneration | −0.34 | −0.62 | 0.010 |
| NR4A2 (204621_s_at) | dopaminergic proxy | −1.08 | −0.56 | 0.023 |
| SNCA | Synuclein–mitochondrial; Proteostasis–autophagy; Global neurodegeneration | −0.63 | −0.53 | 0.032 |
| LRRK2 | Synuclein–mitochondrial; Proteostasis–autophagy | −0.31 | −0.44 | 0.074 |
| NFKB1 | Inflammatory microglial; Astrocyte stress; Immune antigen presentation; Vascular barrier | 0.18 | 0.40 | 0.108 |
| MOBP | Myelin–oligodendrocyte; Global neurodegeneration | −0.54 | −0.40 | 0.108 |
| LAMP5 | Neuronal vulnerability | −0.80 | −0.38 | 0.136 |

*(continued)*

| Gene (probe) | Role | $\log_2$ FC (PD − control) | Rank-biserial | $p$ (exact) |
|---|---|---|---|---|
| GPNMB | Inflammatory microglial; Proteostasis–autophagy; Global neurodegeneration | 0.46 | 0.36 | 0.152 |
| IL1B | Inflammatory microglial; Astrocyte stress; Vascular barrier | 0.42 | 0.36 | 0.152 |
| APOE | Astrocyte stress; Synuclein–mitochondrial; Amyloid–tau; Global neurodegeneration | −0.27 | −0.36 | 0.152 |
| MBP | Myelin–oligodendrocyte; Global neurodegeneration | −0.21 | −0.32 | 0.207 |
| TYROBP | Inflammatory microglial; Immune antigen presentation; Global neurodegeneration | 0.29 | 0.31 | 0.229 |
| KCNJ6 (207446_at) | dopaminergic sensitivity | −0.55 | −0.29 | 0.251 |
| NEFL | Neuronal vulnerability; Global neurodegeneration | −0.73 | −0.29 | 0.251 |
| PSEN1 | Amyloid–tau | −0.27 | −0.28 | 0.276 |
| TREM2 | Inflammatory microglial; Global neurodegeneration | 0.30 | 0.26 | 0.301 |
| AIF1 | Inflammatory microglial; Immune antigen presentation; Global neurodegeneration | 0.56 | 0.26 | 0.301 |
| P2RY12 | Inflammatory microglial | 0.25 | 0.24 | 0.357 |
| NEFM | Neuronal vulnerability; Global neurodegeneration | −0.29 | −0.19 | 0.452 |
| PVALB | Neuronal vulnerability | 0.49 | 0.19 | 0.452 |
| MAPT | Synuclein–mitochondrial; Amyloid–tau; Proteostasis–autophagy; Global neurodegeneration | 0.11 | 0.15 | 0.559 |
| CX3CR1 | Inflammatory microglial; Vascular barrier | 0.22 | 0.14 | 0.598 |
| APP | Amyloid–tau; Proteostasis–autophagy; Global neurodegeneration | 0.05 | 0.11 | 0.677 |
| TNF | Inflammatory microglial; Astrocyte stress; Vascular barrier | −0.14 | −0.10 | 0.718 |
| B2M | Immune antigen presentation; Vascular barrier | 0.08 | 0.10 | 0.718 |
| HLA-DRA | Immune antigen presentation; Global neurodegeneration | 0.02 | 0.08 | 0.760 |
| SLC17A7 | Neuronal vulnerability | 0.05 | 0.06 | 0.846 |
| SLC6A3 (206868_at) | dopaminergic sensitivity | 0.00 | 0.04 | 0.890 |
| GFAP | Astrocyte stress; Global neurodegeneration | −0.01 | −0.01 | 0.978 |
| PLP1 | Myelin–oligodendrocyte; Global neurodegeneration | 0.01 | 0.01 | 0.978 |
| SST | Neuronal vulnerability | 0.08 | 0.01 | 0.978 |

**Table S6. GSE7621 competitive null** (10,000 expression-matched random probe sets per axis; per-gene $z$-score).

| Axis | Observed | Null median | Null 2.5% | Null 97.5% | Percentile | Empirical $p$ |
|---|---|---|---|---|---|---|
| Inflammatory microglial | 0.32 | 0.07 | −0.60 | 0.68 | 76.9 | 0.506 |
| Astrocyte stress | 0.14 | 0.03 | −0.61 | 0.67 | 63.1 | 0.760 |
| Myelin–oligodendrocyte | −0.32 | 0.00 | −0.54 | 0.57 | 15.1 | 0.305 |
| Neuronal vulnerability | −0.18 | 0.01 | −0.58 | 0.67 | 30.5 | 0.605 |
| Synuclein–mitochondrial | −0.60 | 0.00 | −0.58 | 0.65 | 2.4 | 0.062 |
| Amyloid–tau | −0.53 | 0.00 | −0.57 | 0.62 | 4.0 | 0.092 |
| Immune antigen presentation | 0.24 | 0.04 | −0.58 | 0.65 | 72.0 | 0.595 |
| Vascular barrier | 0.31 | 0.03 | −0.60 | 0.65 | 78.4 | 0.439 |
| Proteostasis–autophagy | −0.29 | 0.00 | −0.58 | 0.65 | 19.4 | 0.416 |
| Global neurodegeneration | −0.38 | 0.00 | −0.51 | 0.60 | 8.9 | 0.215 |

**Table S7. GSE7621 composition models for every axis** (per-gene $z$-score; OLS; PD coefficient in SD units).

| Axis | PD only | + sex | + dopaminergic | + dopaminergic + sex | + PC1, PC2 | $\rho$ with proxy |
|---|---|---|---|---|---|---|
| Inflammatory microglial | 0.51 ($p$ 0.224) | 0.54 ($p$ 0.250) | 1.01 ($p$ 0.074) | 1.02 ($p$ 0.089) | 0.80 ($p$ 0.109) | −0.04 |
| Astrocyte stress | −0.04 ($p$ 0.919) | 0.29 ($p$ 0.502) | 0.06 ($p$ 0.925) | 0.34 ($p$ 0.547) | −0.10 ($p$ 0.814) | −0.03 |
| Myelin–oligodendrocyte | −0.50 ($p$ 0.241) | −0.41 ($p$ 0.376) | −1.37 ($p$ 0.010) | −1.27 ($p$ 0.021) | −0.93 ($p$ 0.041) | −0.14 |
| Neuronal vulnerability | −0.26 ($p$ 0.541) | −0.47 ($p$ 0.311) | 0.75 ($p$ 0.132) | 0.54 ($p$ 0.265) | 0.09 ($p$ 0.788) | 0.44 |
| Synuclein–mitochondrial | −1.13 ($p$ 0.004) | −1.26 ($p$ 0.003) | −0.27 ($p$ 0.507) | −0.40 ($p$ 0.329) | −0.85 ($p$ 0.039) | 0.56 |
| Amyloid–tau | −1.05 ($p$ 0.009) | −1.04 ($p$ 0.017) | −1.23 ($p$ 0.021) | −1.22 ($p$ 0.030) | −1.14 ($p$ 0.019) | 0.18 |
| Immune antigen presentation | 0.41 ($p$ 0.334) | 0.59 ($p$ 0.203) | 0.88 ($p$ 0.124) | 1.02 ($p$ 0.087) | 0.53 ($p$ 0.292) | 0.01 |
| Vascular barrier | 0.36 ($p$ 0.403) | 0.49 ($p$ 0.293) | 0.61 ($p$ 0.291) | 0.72 ($p$ 0.236) | 0.26 ($p$ 0.607) | 0.05 |
| Proteostasis–autophagy | −0.64 ($p$ 0.124) | −0.86 ($p$ 0.057) | 0.11 ($p$ 0.819) | −0.09 ($p$ 0.852) | −0.40 ($p$ 0.354) | 0.35 |
| Global neurodegeneration | −0.56 ($p$ 0.183) | −0.48 ($p$ 0.294) | 0.18 ($p$ 0.722) | 0.23 ($p$ 0.670) | −0.23 ($p$ 0.631) | 0.44 |

Dopaminergic proxy: TH 208291_s_at, SLC18A2 205857_at, DDC 205311_at, ALDH1A1 212224_at, NR4A2 204621_s_at, RET 211421_s_at (mean probe $z$-score). Median pairwise Spearman $\rho$ among the six probes 0.77. Sensitivity proxy adds SLC6A3 206868_at and KCNJ6 207446_at.

**Table S8. Simulation summary** (200 cohorts per scenario; proportion of cohorts in each class).

| Scenario | Axis | Naive significant | Composition-independent | Composition-explained | Median rank-biserial |
|---|---|---|---|---|---|
| No effect | Inflammatory microglial | 0.035 | 0.035 | 0.000 | 0.01 |
| No effect | Neuronal vulnerability | 0.005 | 0.005 | 0.000 | −0.02 |
| No effect | Synuclein–mitochondrial | 0.005 | 0.005 | 0.000 | −0.01 |
| Neuron loss only | Inflammatory microglial | 0.005 | 0.000 | 0.005 | 0.00 |
| Neuron loss only | Neuronal vulnerability | 0.815 | 0.050 | 0.765 | −0.61 |
| Neuron loss only | Synuclein–mitochondrial | 0.020 | 0.010 | 0.010 | 0.01 |
| Neuron loss + weak direct | Inflammatory microglial | 0.755 | 0.495 | 0.245 | 0.52 |
| Neuron loss + weak direct | Neuronal vulnerability | 0.875 | 0.035 | 0.840 | −0.59 |
| Neuron loss + weak direct | Synuclein–mitochondrial | 0.025 | 0.005 | 0.020 | −0.01 |
| Neuron loss + moderate direct | Inflammatory microglial | 1.000 | 0.975 | 0.025 | 0.81 |
| Neuron loss + moderate direct | Neuronal vulnerability | 0.930 | 0.030 | 0.900 | −0.58 |
| Neuron loss + moderate direct | Synuclein–mitochondrial | 0.015 | 0.000 | 0.015 | 0.00 |
| Moderate direct only | Inflammatory microglial | 1.000 | 1.000 | 0.000 | 0.80 |
| Moderate direct only | Neuronal vulnerability | 0.010 | 0.000 | 0.010 | −0.01 |
| Moderate direct only | Synuclein–mitochondrial | 0.025 | 0.015 | 0.010 | −0.03 |

### Supplementary Figures

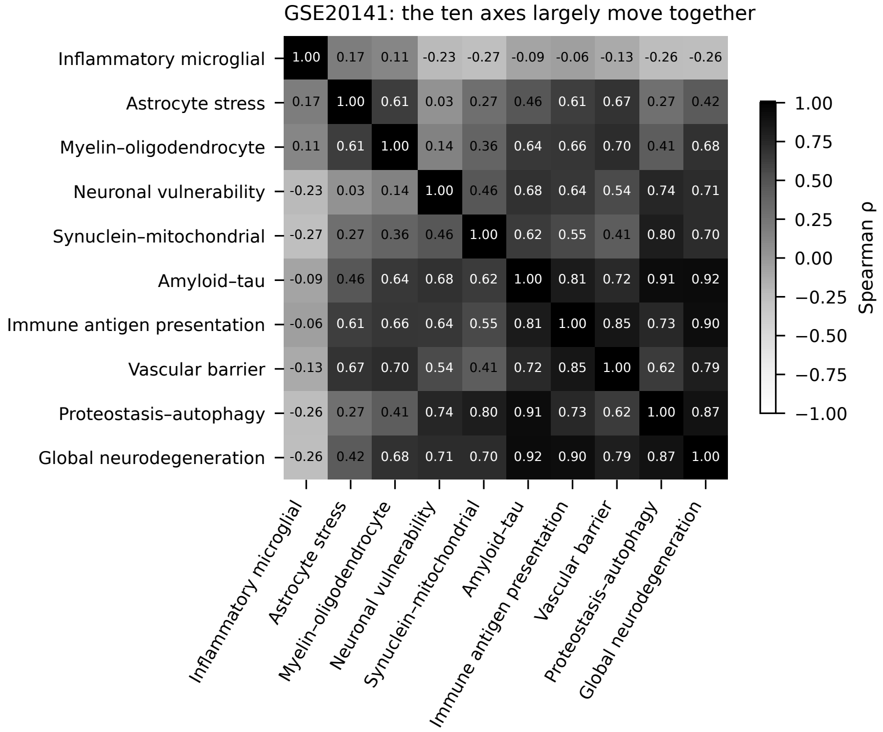


**Figure S1. GSE20141 axis inter-correlation.** Spearman correlations between the ten axis scores written by the original pipeline (18 samples). The nominally distinct axes largely move together.

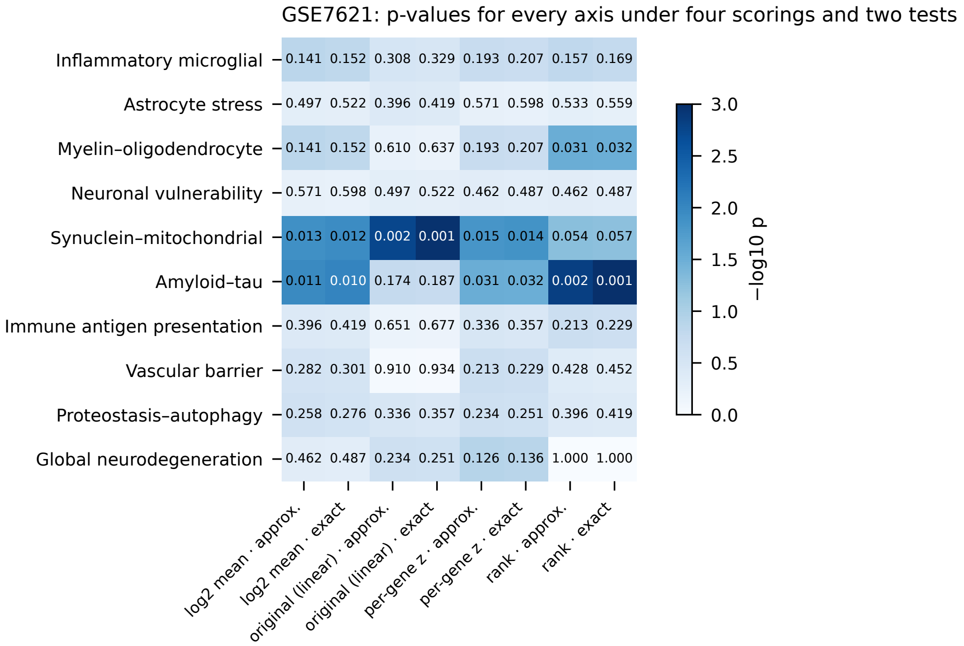


**Figure S2. GSE7621 specification grid.** $P$-values for every axis under four scorings (original linear, $\log_2$ mean, per-gene $z$-score, rank) and two tests (normal approximation, exact). Which axes reach $p < 0.05$ depends on the scoring.

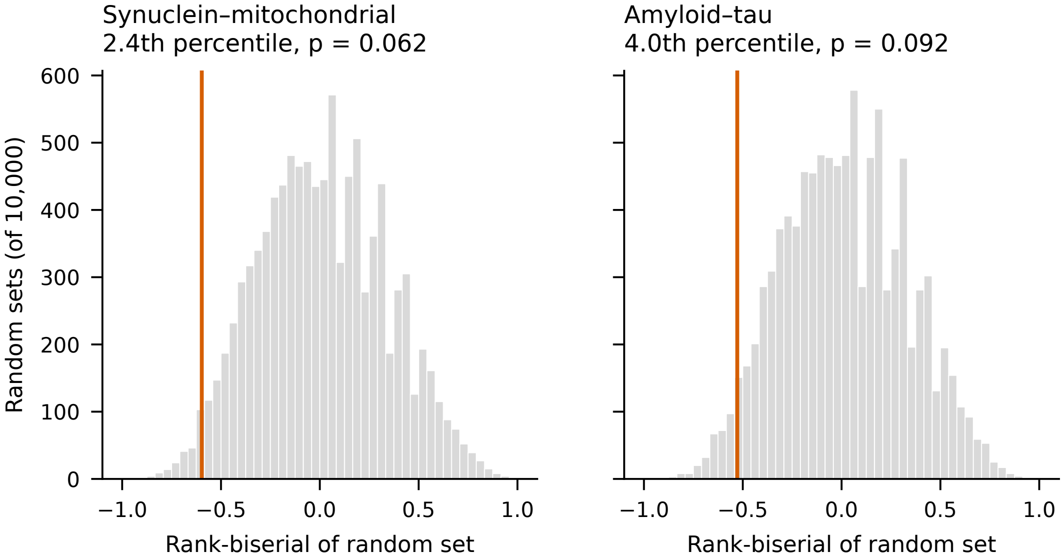


**Figure S3. Competitive null distributions in GSE7621.** Rank-biserial effects of 10,000 expression-matched random probe sets for the two axes with the lowest *p*-values; red line, observed axis.

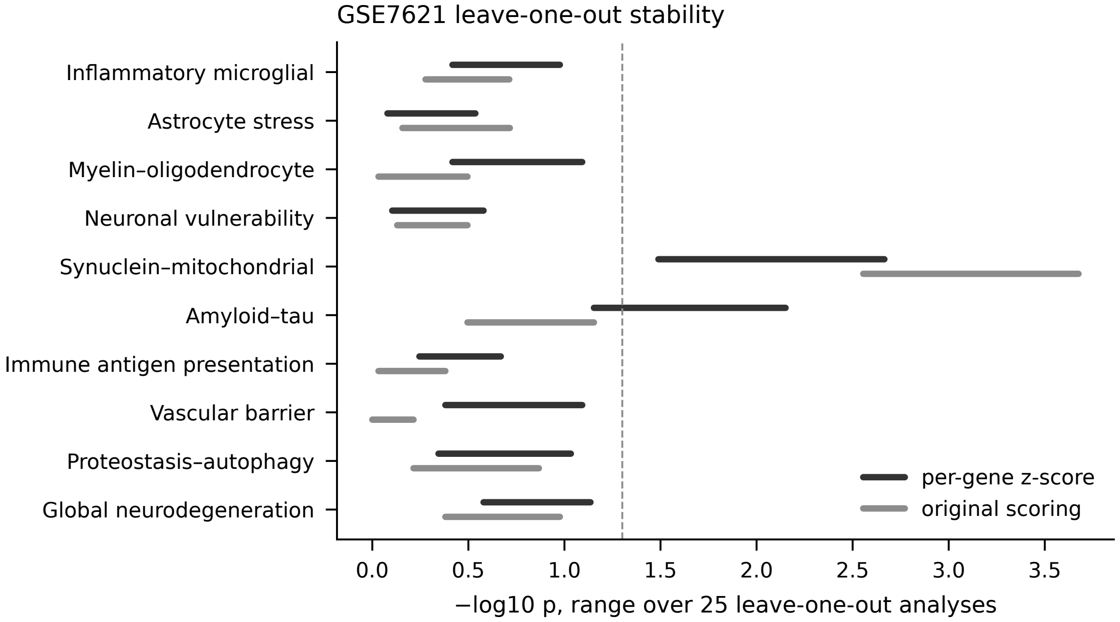


**Figure S4. Leave-one-out stability in GSE7621.** Range of $-\log_{10}$ exact $p$ over the 25 leave-one-out analyses, for the per-gene $z$-score and the original score; dashed line, $p = 0.05$.

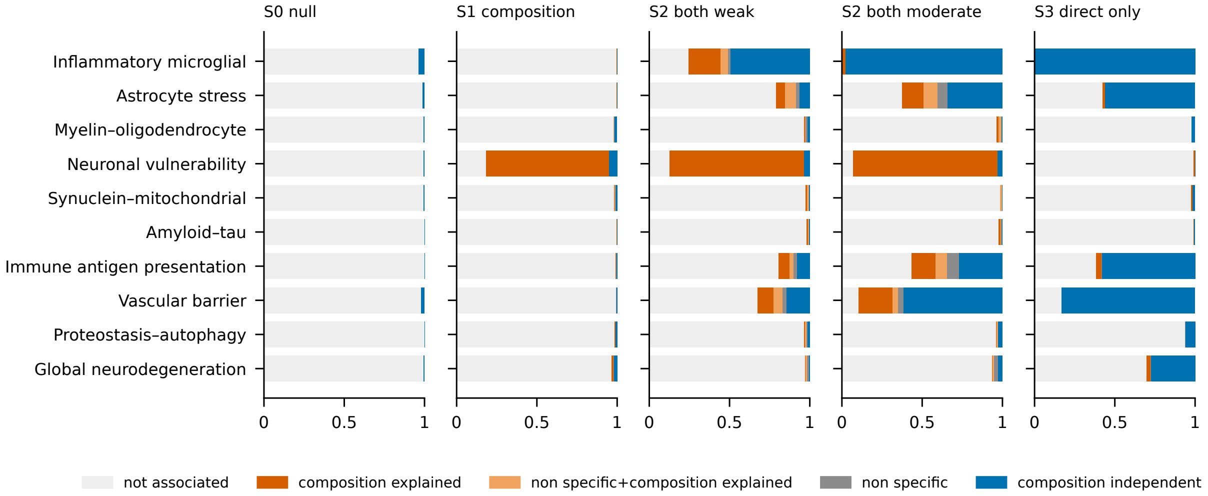


**Figure S5. Battery classes for every axis in every simulated scenario.** Gene sets that share genes with the planted set (astrocyte stress, immune antigen presentation, vascular barrier, global) are detected as composition-independent when the planted effect is present, as expected.

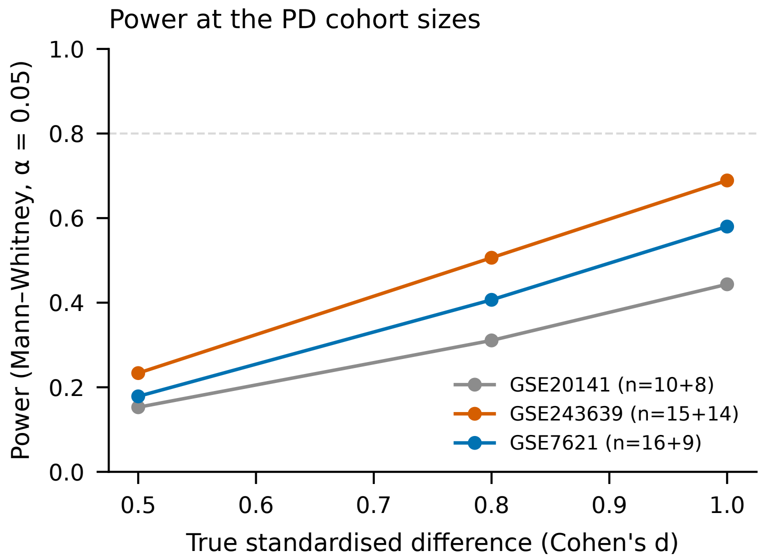


**Figure S6. Power of the two-sided Mann–Whitney test at the PD cohort sizes**, estimated from 4,000 simulated normal samples per point.
